# Priming using Human and Chimpanzee Expressions of Emotion Biases Attention toward Positive Emotions

**DOI:** 10.1101/2023.10.20.562671

**Authors:** Anna Matsulevits, Mariska E. Kret

## Abstract

Perceiving and correctly interpreting emotional expressions is one of the most important abilities for social animals’ communication. It determines the majority of social interactions, group dynamics, and cooperation, being highly relevant for an individual’s survival. Core mechanisms of this ability have been hypothesized to be shared across closely related species with phylogenetic similarities. Here, we explored homologies in human processing of different species’ facial expressions using eye-tracking. Introducing a prime-target paradigm, we tested the influences on human attention elicited by priming with differently valenced emotional stimuli depicting human and chimpanzee faces. We demonstrated an attention shift towards the conspecific (human) target picture that was congruent with the valence depicted in the primer picture. We did not find this effect with heterospecific (chimpanzee) primers and ruled out that this was due to participants interpreting them incorrectly. Implications about the involvement of related emotion-processing mechanisms for human and chimpanzee facial expressions, are discussed. Systematic cross-species-investigations of emotional expressions are needed to unravel how emotion representation mechanisms can extend to process other species’ faces. Through such studies, we address the gap of a shared evolutionary ancestry between humans and other animals to ultimately answer the question of *“Where do emotions come from?”*.

## Introduction

In social animal groups, the ability to integrate different cues from facial expressions largely determines inter-individual interactions and influences group dynamics and cooperation, conceivably having an impact on survival (Bourjade, 2016). Nevertheless, the mechanisms by which our brains process these sensory cues from species other than humans remain relatively unexplored. In the current study, we aspired to investigate the underpinnings of the discernment of complex facial muscle movements – manifestations of emotional expressions. We tested whether humans possess the capacity to interpret and comprehend the facial expressions of one of our genetically closest kin, the chimpanzee (*Pan troglodytes*), and to what extent their priming influence endures, as compared to emotional expressions within our own species.

While the importance of emotions in regulating and navigating the lives of social species is well established (van Hoof, 1967), the universality of emotional expressions, their contagion, and how we and other animals process them remain controversial. Referring to Darwin’s theory of emotion universality (1872), humans and other animals show emotional states through remarkably similar face-and-body actions. Similar adaptive properties that apply across species further support cognitive underpinnings of emotion processing being well conserved among humans and other great apes (Anderson & Adolphs, 2014; Nieuwburg, Ploeger & Kret, 2021). To effectively process such signals, it is required to form an internal representation of the other’s emotion based on the information from different facial muscle movements and their complex combinations (Nieuwburg et al., 2021). As a result of their nearly identical-to-human facial musculature (Burrows et al., 2006; Waller et al. 2006), chimpanzees are capable of activating a high variety of distinct combinations of so-called action units (AUs) that can resemble even more complex and equivalent-to-human facial expressions (Vick et al., 2007). Supporting the ability of great apes to produce comparable facial expressions, Waller, Cray, and Burrows (2008) found facial muscles which are involved in the production of the basic emotional expressions to show the least individual variation across species.

Whether the internal representation of human emotion can be translated to other species and to what extent we are able to assess emotions in heterospecifics is not entirely clear. Based on the theory of emotion universality and facial expressions having an innate basis leading to a stereotypical appearance, evolutionary closely related species should be able to process each other’s facial expressions (Pollick & de Waal, 2007). If faces of different species that roughly share the spatial arrangement of face elements (eyes, nose, mouth, etc.) are associated with one and the same face prototype, the formation of emotion representations and the correct interpretation of these may be facilitated. Hence, a particular expression in two different species would, in this case, be classified as homologous and assigned to the same category, presumably eliciting similar reactions in the observer. Using the first objective and standardized instrument (ChimpFACS) salient expressions communicating agonistic and affiliative affective states have been validated with a fair degree of certainty in the chimpanzee (Parr et al, 2007; Parr & Waller, 2006). However, comparative studies on visual perception and attention toward affective stimuli yield mixed results. Since it is highly likely that primates’ emotional expressions share a common ancestry, it is not surprising that humans can differentiate between facial expressions of their own, and expressions of other primates (Kret, Muramatsu & Matsuzawa, 2018; Parr, 2003). For example, affective stimuli depicting heterospecifics has shown to induce similar interpretation and attention bias, evoking comparable physiological reactions as equivalent stimuli depicting conspecifics (Dufour, Pascalis & Petit, 2006; Parr, Hopkins & de Waal, 1998; Parr, 2001; Berlo, 2022).

On the other hand, multiple studies in human and non-human primates have confirmed better processing and understanding of information observed in a conspecific (Hattori, Kano & Tomonaga 2010). Thus, it remains unclear to what extent affective states in other, evolutionary closely related but not domesticated species can be processed, and in which way those exert influence on one’s attention, perception, and potentially one’s emotional state.

Being generally more attentive towards affectively significant stimuli of evolutionary importance is highly beneficial since quickly reorientating visual attention and correctly evaluating such signals increases the chances of survival (Vuilleumier, 2005). Regardless of age, species, and the ability to consciously process and understand an affective expression, attentional biases towards emotional stimuli are overall a well-studied concept (van Rooijen, Ploeger & Kret, 2017). In natural environments, especially in potentially threatening situations, spotting negative cues is more important to make a quick decision about the subsequent action, namely to *fight* or *flight* (Frijda, 2010). Therefore, attentional biases are often observed towards cues with a negative association (Vaish, Grossmann & Woodward, 2008). Given their evolutionary role, threatening faces in visual search studies are detected faster with an increasing perception of “threat” (Öhman, Lundqvist & Esteves, 2001; Wilson & Tomonaga; 2021). Contradictory to this finding, Smith et al. (2006) argued that in environments where the benefits from positive events are outweighing the consequences from negative events, individuals might adapt to be more prone toward positive rather than negative information. This adaptation seems to be bidirectional (Aas er al., 2017; Dai et al., 2016). Put more generally, such shifts in attention towards particular valenced affective stimuli could potentially be partly traced back to the accumulated influences which are prevalent in an agent’s environment.

By applying semantic prime-target paradigms, the effects of prior context on the perception of a subsequent stimulus can be used to study the recognition and processing of emotional expressions (Carroll & Young, 2005). Presenting a cue that is related to a subsequently shown target has been found to change the attention, reaction, and classification accuracy toward that target (Higgins, Bargh & Lombardi, 1958). Once a priming stimulus is perceived, it is thought to trigger a cascade of associations in the receiver, facilitating and fastening the processing of related stimuli (Molden, 2014). Demonstrating this effect, Smith et al. (2006) found attentional bias in participants who were primed with positive and negative information to shift toward a stimulus that was congruent with the primer. Semantic priming has consistently been demonstrated to increase reaction times for non-matching targets and decrease reaction times for matching targets (McNamara, 2005). This further indicates a tendency of adaptation and modulation of bias based on the demands of an individual’s situation or environment (Smith et al., 2006), supporting the idea of Fredrickson’s (1998) broaden-and-build model. In his work, Fredrickson suggests that positive emotions would temporarily broaden an individual’s thought-action repertoire, promoting attention to related (here: positively valenced) cues. This theory was confirmed by Wadlinger and Isaacowitz (2006), who found positive mood (induced by sugary food) to broaden visual attention to positive stimuli. Carroll and Young (2005) conducted several studies on affective priming with facial expressions, showing that facial expressions can likewise act as direct elicitors of affect, which is in line with facial displays being seen as tools for social influence (Crivelli & Fridlund, 2018). Interestingly, Carroll and Young (2005) demonstrated that this priming effect on recognition can cross from nonverbal representations (pictures of emotions) to verbal representations (words) and vice versa. Given the discussed parallels between chimpanzee and human faces, along with the assumption of the ability to correctly classify facial expressions in heterospecifics, this bidirectional priming effect could apply to primers depicting different species’ facial expressions.

Comparing how differently valenced emotional primers of different species influence the attention toward emotional target stimuli will further improve our understanding of the mechanisms underlying the internal representations of affective processing. If primers of different species are understood correctly, then they would identically facilitate the accessibility of positive and negative constructs in memory, depending on the valence category of the primer, irrespective of the species. By triggering associated perceptual representations, an affective primer could potentially shift the observer’s mood, introducing biased perception towards stimuli related to the induced mood. Therefore, the perceived primer valence might influence the agent’s own internal state to the extent that it shifts the attentional biases towards a matching state. Supporting this assumption, previous work has confirmed mood to modulate attentional biases (Rowe, Hirsch & Anderson, 2007). We aim to explore the functional consequences of facial expressions and broaden our knowledge of how such affective stimuli of different species are understood and how they impact attention-related processes in the viewer.

## Hypotheses

**H_1_:** Priming with emotionally valenced faces (positive and negative), compared to neutral faces of both species (i.e., humans and chimpanzees) introduces an attentional bias towards the emotion representations on the target screen.

**H_2_:** The direction of the attention shift depends on the congruency with the primer valence. Attention is shifted towards the stimulus on the target screen that is congruent with the valence previously presented in the primer picture.

**H_3_:** The elicited attentional bias is larger upon seeing pictures of conspecifics compared to seeing heterospecifics.

## Methods

### Participants

A total of 50 participants recruited at Leiden University took part in the eye-tracking experiment after filling in the informed consent. The participants were reimbursed with course credit (1 credit for 30 minutes of participation). The sample consisted of 30 women and 20 men with an average age of 26.5 years old (*SD* = 6.59). All participants had normal or corrected-to-normal vision and no history of clinically diagnosed psychiatric or neurological conditions. Data were collected in June 2022. The procedure and methods were approved by the Leiden University Ethics Committee (CEP: 2022-02-20-M.E. Kret-V1-3988).

### Stimulus material

The human face stimuli were taken from the validated Chicago Face Database (CFD), while the chimpanzee face stimuli were collected from different resources such as researcher’s archives, animal photographers’ work, and the iNaturalist webpage for uploading high quality pictures of different species, suitable for research purposes. The stimuli set that was used for the primer pictures contained 18 unique primer pictures for each of the six conditions, resulting in 108 primer pictures in total. No primer pictures were re-used as target stimuli.

Since each trial required an affiliative and an agonistic picture of a human for the target screen, our stimulus set for the targets consisted of (108 trials * 2 valences) 216 target pictures in total. We could have had 216 unique target pictures depicting human emotional expressions, however, due to the limited resources of emotional stimuli of chimpanzees, we were not able to entirely avoid repetition of the emotionally valenced target pictures in this group. The additional material was also taken from the Chicago Face Database. Thus, we added 27 extra pictures of positively and 27 extra pictures of negatively valenced human emotional expressions to our stimulus set which were repeated 4 times during the trials (27 pictures * 2 valences * 4 repetitions = 216). To account for this limitation, we made sure that 1) there is no overlap between the primer and one of the target pictures within the same trial 2) the target does not contain any picture from the previous and the next trial, and 3) the position of the target (left/right) regarding the emotional valence depicted, is pseudo-randomized and counterbalanced across trials and sessions. The colored pictures had a dimension of 420 x 320 pixels on a 1280 x 1024 computer display.

We selected stimuli of emotional expressions in humans and chimpanzees that appear to be well represented across these species. For the affiliative (positive) pictures of humans, we selected images that were labelled as “happy” in the CFD and for the agonistic (negative) pictures, we selected images that were labelled as “angry”. These emotional categories are found in other primates and equivalent facial expressions communicating these internal states have been observed in chimpanzees. Expressing an angry face for humans includes the activation of AU4, AU7, AU10, AU16, AU25, AU5, AU20, AU9 and AU26, whereas expressing a happy face includes the activation of AU12, AU7, AU26, AU6, AU10, AU1 and AU25 (Kohler et al., 2004). Hence, there are only three overlapping AUs among these human expressions, making them good models for studying emotional function. Nevertheless, relying purely on the AU activation for deducing similarities in expressions between species can be misleading. Entangling the activation of facial muscles in chimpanzees for expression production has shown that some identical AUs (i.e., AU10, AU16) were indeed active for chimpanzee agonistic faces as for human angry and fearful (agonistic) faces. However, finding homologous expressions in the prototypical chimpanzee facial expression repertoire is more challenging, because a related expression can communicate different affective signals. For instance, the AUs active in a human smile are overlapping with AUs in fear-grin and bared-teeth displays in chimpanzees (i.e., AU12, AU25) which occur predominantly in stressful situations (Parr & Waller, 2006). This makes two facial expressions communicating contrasting signals in humans and chimpanzees, related to each other.

For the equivalent of a “happy” face in chimpanzees we chose to base our affiliative stimuli selection following the proposed parallels between human laughter and a non-human “play-face” (van Hooff, 1972). All AUs present in a chimpanzee play-face (AU12, AU25, AU26) are within the subset of a human happy-face (Parr &Waller, 2006). For the selection of agonistic stimuli, we matched the “angry” face in humans with the bared-teeth and screaming displays in chimpanzees. Knowing that this could potentially be a source for interpretation mistakes in our participants and consequently lead to wrong priming effects, we compared their classification ratings (in valence and arousal of the seen stimuli) with the ratings of seven non-human primate experts and found no significant difference between the non-experts and experts (see Supplementary Material I. Validation of the Stimulus Material). In addition, we confirmed that valence and arousal ratings of human and chimpanzee stimuli were in line with our expectations (see Supplementary Material Figure 2).

### Calibration

Participants were calibrated using the 5-point automated calibration procedure in Tobii Pro Lab. Calibrations were accepted when the error displayed after finishing the calibration was minimal (less than one degree) and the data loss was less than 1%.

### Design and Procedure

Participants were actively recruited by the experimenter in the facilities of Leiden University. After reading the information letter for the study and signing the consent form, participants were individually tested in an eye-tracking lab at Leiden University. By signing the consent form they allowed to use their data for further analyses and publications.

We developed a within-subject design with three predictor conditions (affiliative, agonistic, neutral) of the priming factor and two valence levels (affiliative, agonistic) shown on the target screen. In the prime-target paradigm, positively and negatively valenced picture targets of emotional expressions were presented adjacently (4 seconds), preceded by a positive,negative, or neutral facial expression prime of either a human or chimpanzee face (2 seconds). Importantly, the target screen always depicted affiliative and agonistic human facial expressions. The order of the trials and the position of the positive and negative target images (left/right) were pseudo-randomized and counterbalanced across trials and sessions. Two trial sequences of different *primer species* conditions can be found in Figure 1.

**Figure 1.**
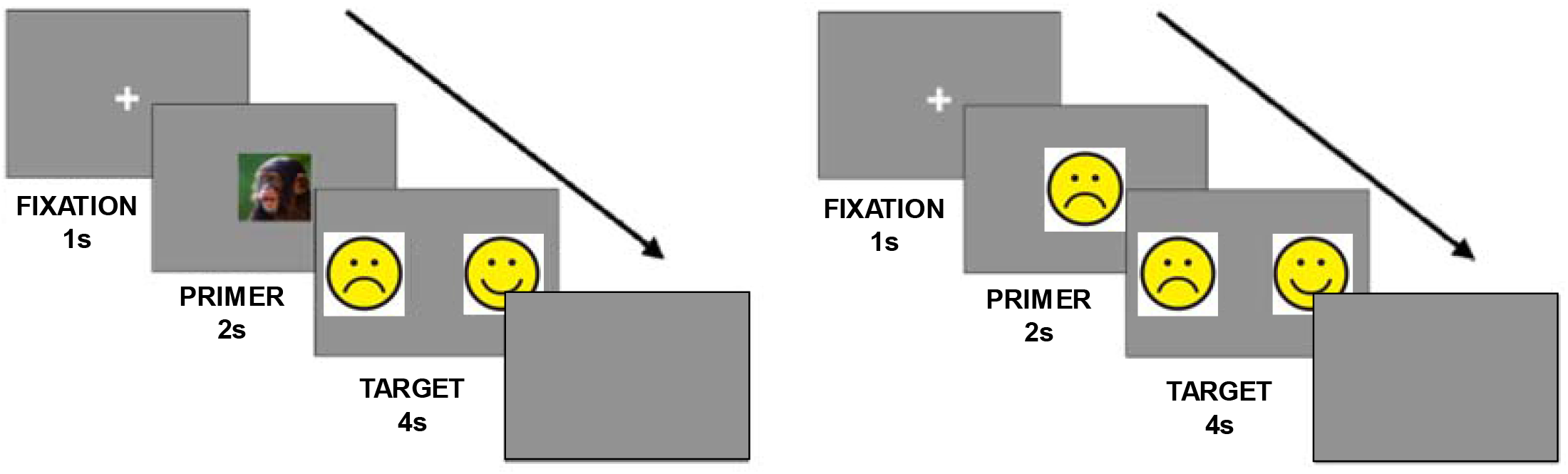
*Note.* Trial sequence. Each trial started with a fixation cross (1s), followed by the first screen with the stimulus of a primer (2s) and subsequently by the second screen with the two target stimuli presented next to each other (4s). The trial ended with a blank screen (3s). *** The real stimuli photographs of human faces are replaced with emoji faces only for the present preprint version in order to meet the policies of *biorxiv*. ***

The experiment was run via Tobi Pro Lab (version 1.181.37603) on a Windows computer. After the participant sat down behind the computer, the experimenter started the 5-point calibration procedure (duration ∼ 1 minute). Once the calibration was finished, the program proceeded to the eye-tracking trials. During the eye-tracking procedure, the participants had no active task to perform, but were asked to freely view the images on the computer screen. Their eye movements were measured via a Tobii Pro-Fusion eye-tracker attached to the monitor. A session started with a fixation cross that was shown for 1000ms. This was followed by a primer depicting the facial expression of a human or a chimpanzee (classified as either affiliative, agonistic, or neutral) that flashed up for 2000ms in the middle of the screen, then directly followed by a 4000ms target screen showing two emotionally valenced pictures depicting humans. The primer picture was spatially not overlapping with the position of the target pictures to prevent inaccuracies in the gaze fixation assessment. A trial ended with a blank screen that was shown for 3000ms. After the 9^th^ session, a short break was programmed into the experiment, so that the participants could rest their eyes and look away for a couple of seconds. Once the second part was finished (108 trials in total), a Qualtrics questionnaire was opened remotely on the participant’s screen that first assessed the participants’ demographic information and then proceeded to the rating task. Participants were asked to rate all the primer pictures (108 in total) on two separate sliders, both on valence (negative to positive) and arousal (low to high). Since the primer pictures were only shown once, keeping the familiarization effects at its minimum compared to the target pictures, we decided to limit the rating to the sub-set of the primer pictures. The answers were coded on a 100-point scale with 50 indicating the neutral “zero-point” of the slider. Numbers smaller than 50 represented the rating in the negative/low spectrum and numbers larger than 50 represented the rating in the positive/high spectrum. Participants were given a debrief form explaining the background information and the goal of the study after they had finished the task, as well as the opportunity to ask remaining questions. The experiment took about 30-40 minutes to finish.

### Data Preparation

Before the analyses, we plotted the gaze data with the locations of the stimuli on the screen to check whether the raw fixation data matched with the areas of the stimuli on the screen. We drew a 430×320 square around each of the primer pictures and around each of the two simultaneously presented target pictures. Through Tobii Pro Lab’s *Metrics* option, we extracted the data on *Total Fixation Duration* per ROI using the Tobii Pro Lab Fixation Filter.

### Statistical Analyses

All data was analyzed in RStudio (v. 4.1.2). To answer our research questions and test whether emotional primers elicit attentional bias, we performed a multi-level analysis using Bayesian mixed modeling to analyse the total fixation duration. Our key question was whether fixations on the emotionally valenced targets were influenced by the previously seen primer emotion and species. Since the target screen depicted two facial expressions simultaneously, the looking durations toward the targets were highly correlated. Thus, we calculated our dependent variable from the proportional looking duration towards the positive target picture (based on Tobi Pro Lab’s *Total Fixation Duration* (TFD), from here on: bias score) using the following formula:

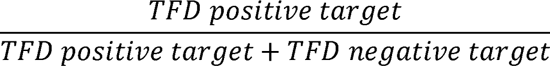

Since the bias score reflects the probability of looking at the positive picture, the “remaining” probability is the attentional bias towards the negative picture. Thus, there is no need of computing an extra negative bias score. The measure of the bias score higher than 0.5 indicates a longer fixation duration towards the positive emotional expression in the target screen. Hence, a bias score lower than 0.5 indicates a longer fixation duration towards the negative emotional expression.

To analyze the eye-tracking data, we used a zero-one inflated Bayesian beta regression model, which is suitable to analyze continuous proportions containing zeros and ones. These models consist of two components, namely a beta component to describe the values between 0 and 1, and a binary component to predict the occurrences of zeros and ones (Ospina & Ferrari, 2021). For our measure of interest, proportional looking duration to emotional target stimuli (positivity bias) across trials, we ran a multilevel model analyzing the main effects, as well as the interaction between variables *primer species (2)* and *primer emotion valence (3)*. Since the dependency between scores from the same participant violates a key assumption of the linear regression, using a multilevel model was best suited to account for these correlations in the data. All variables of interest were dummy-coded and modeled as fixed effects. We allowed the intercept to vary by subject. Then, we computed the positivity bias and assessed whether these differed depending on emotion (agonistic, affiliative, or neutral primer) and the species (human or chimpanzee primer), or their interaction.

For the independent variables, we specified regularizing Gaussian priors with M = 0 and SD = 0.5. This also applied to the independent variables in the formulas for phi, coi, and zoi. For all variance parameters, we kept the default Student’s t priors with 3 degrees of freedom. Furthermore, we kept the default logistic priors for the intercepts of zoi and coi, and default Student’s t prior with 3 degrees of freedom for the Intercept of phi. After running the model, we used the emmeans-package (Lenth et al., 2018) to integrate the different model components and provide estimates based on the posterior predictive distribution. Using these values, we calculated multiple quantitative measures to describe the effects. The model notation including prior distributions can be found in the supplementary information.

For the interaction model, we report the medial estimate coefficients, the logit transformed regression coefficients, and the odds ratio coefficients together with the 95% credible interval (CI). In addition, we also report the probability of direction (pd), which indicates the certainty that an effect goes in a specific direction. All analyses were conducted using RStudio (v. 4.1.2) and the packages *brms, emmeans,* and *ez*.

## Results

The results show that firstly, looking duration (positivity bias) on the emotional target pictures is higher than 0.5. This effect was robust for primers depicting humans (*Mdn* = .510, 95% *CI* [0.501 - 0.518], *pd* = 99%), as well as primers depicting chimpanzees (*Mdn* = .515, 95% *CI* [0.506 - 0.523], *pd* = 98%), meaning that all priming effects combined (positive, negative, neutral) led to human participants reliably looking longer at the affiliative compared to agonistic target pictures (see Table 1, Model 1).

**Table 1.**
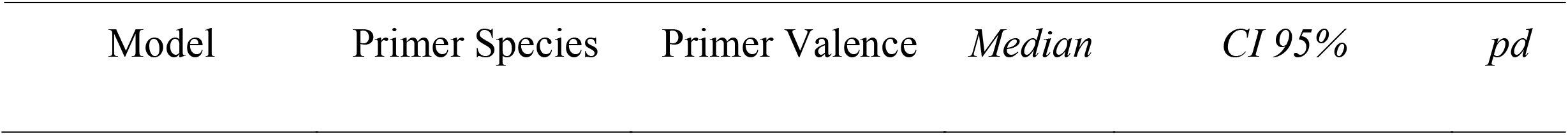
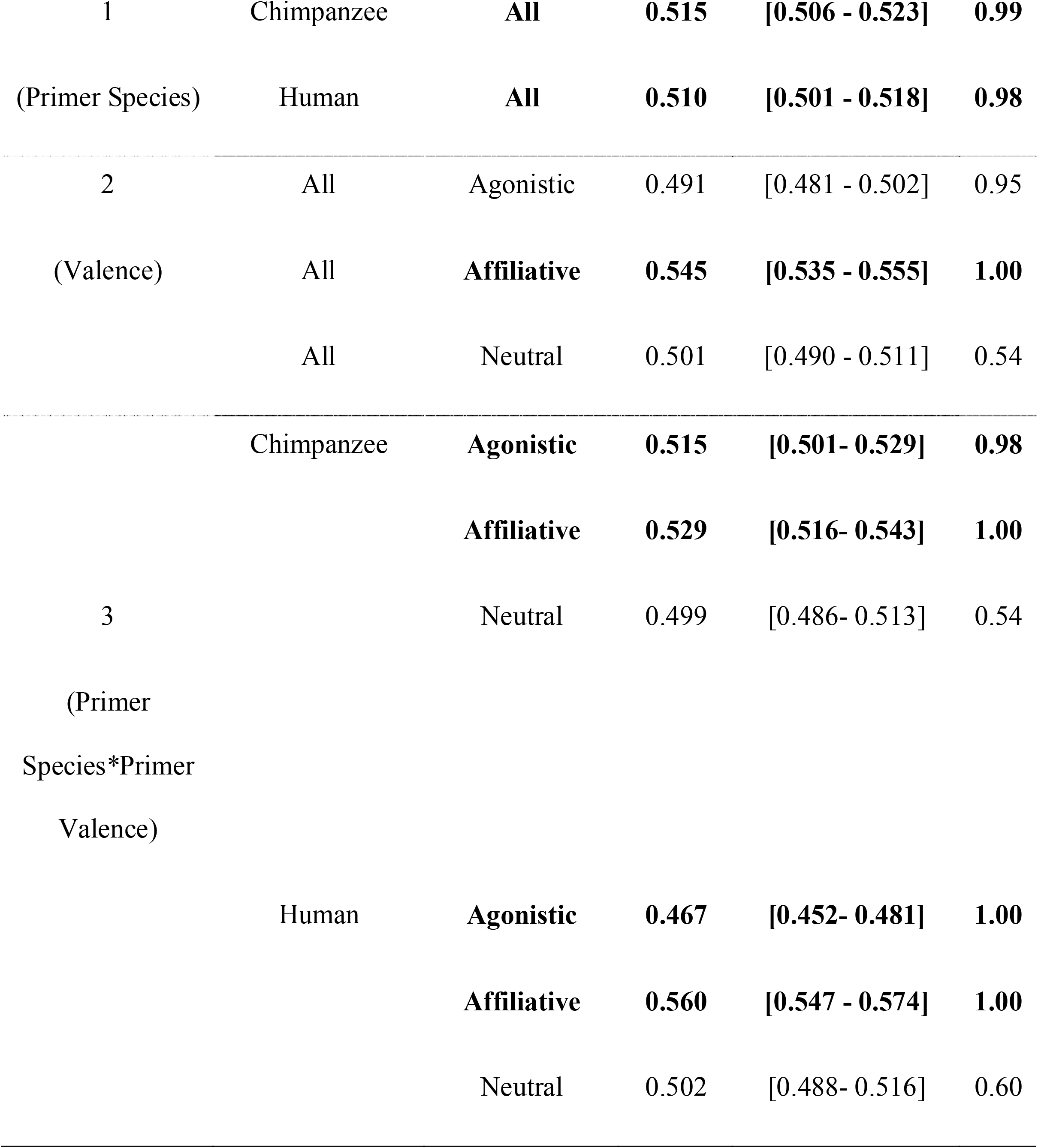
Overview of results per factor level of interest for the three models. Robust effects are in bold.

We then separately investigated the specific emotion categories (*primer valence*), as well as their interaction effect with the *primer species*. In the model where we included *primer valence* as a factor (H_1_), we found robust evidence for increased positivity bias and hence, a decreased negativity bias when the participants were primed with an affiliative primer (*Mdn* = .545, 95% *CI* [0.535 - 0.555], *pd* = 100%) compared to when the primer was either neutral or negative. Summarized across both species, there was no robust effect from seeing agonistic or neutral primers. Hence, we could confirm that averaged over species, only affiliative emotional primers introduce an attentional bias compared to neutral and agonistic primers (H_1_) (see Table 1, Model 2).

Examining the interaction between *primer valence* and *primer species* (H_2,_ H_3_), we observed that attention shift towards the affiliative stimulus on the target screen was linked to presenting chimpanzee primers of affiliative nature (*Mdn* = .471, 95% *CI* [0.457-0.484], *pd*= 1.00), as well as presenting human primers of affiliative nature (*Mdn* = .485, 95% *CI* [0.426 - 0.453], *pd*= 1.00. Furthermore, we found robust evidence for human participants looking longer at the agonistic stimulus in the target screen compared to the affiliative stimuli, given an agonistic human face primer (*Mdn* = .533, 95% *CI* [0.519-0.548], *pd*= 1.00). The opposite effect was found for primers depicting agonistic chimpanzee faces (*Mdn* = .485, 95% *CI* [0.471-0.499], *pd*= 0.98) (H_3_). This disparity also drives the main effect of agonistic valenced primers to being not robust. Neutral primers of both species did not introduce any reliable effect, thus, seeing a neutral primer did not cause a shift in the attention toward a positively or negatively valenced target picture. Entangling this interaction effect confirms that priming with emotionally valenced faces (positive and negative), compared to neutral faces of both species (i.e., humans and chimpanzees) introduces an attentional bias toward the emotion representations in the target screen (H_1_). Inspecting the main effect of valence, this result is not present due to agonistic primers of both species presumably having contrary effects and cancelling each other out.

Zooming in on the interaction effect, we compared the amount of positivity bias introduced by differently valenced emotional primers showing different species (Table 2). In addition we specified the model to estimate precision of the beta distribution, the zero–one inflation probability and the conditional one-inflation probability as a function of the positivity bias. We found the main effect of *primer species* to be significant, with seeing human primers leading to a decreased positivity bias compared to seeing chimpanzee primers (*β*_species_human_= −.20, *CI* [−0.12 – (−0.28)], *OR* = 1.22). For the chimpanzee primers (reference category), there was no significant difference found in positivty bias regardless of the primer valence. This is confirmed by the overlapping credible intervals for the interaction effect between chimpanzee primer and the three valence levels (see Table 1, model 3 and Figure 2).

**Table 2:**
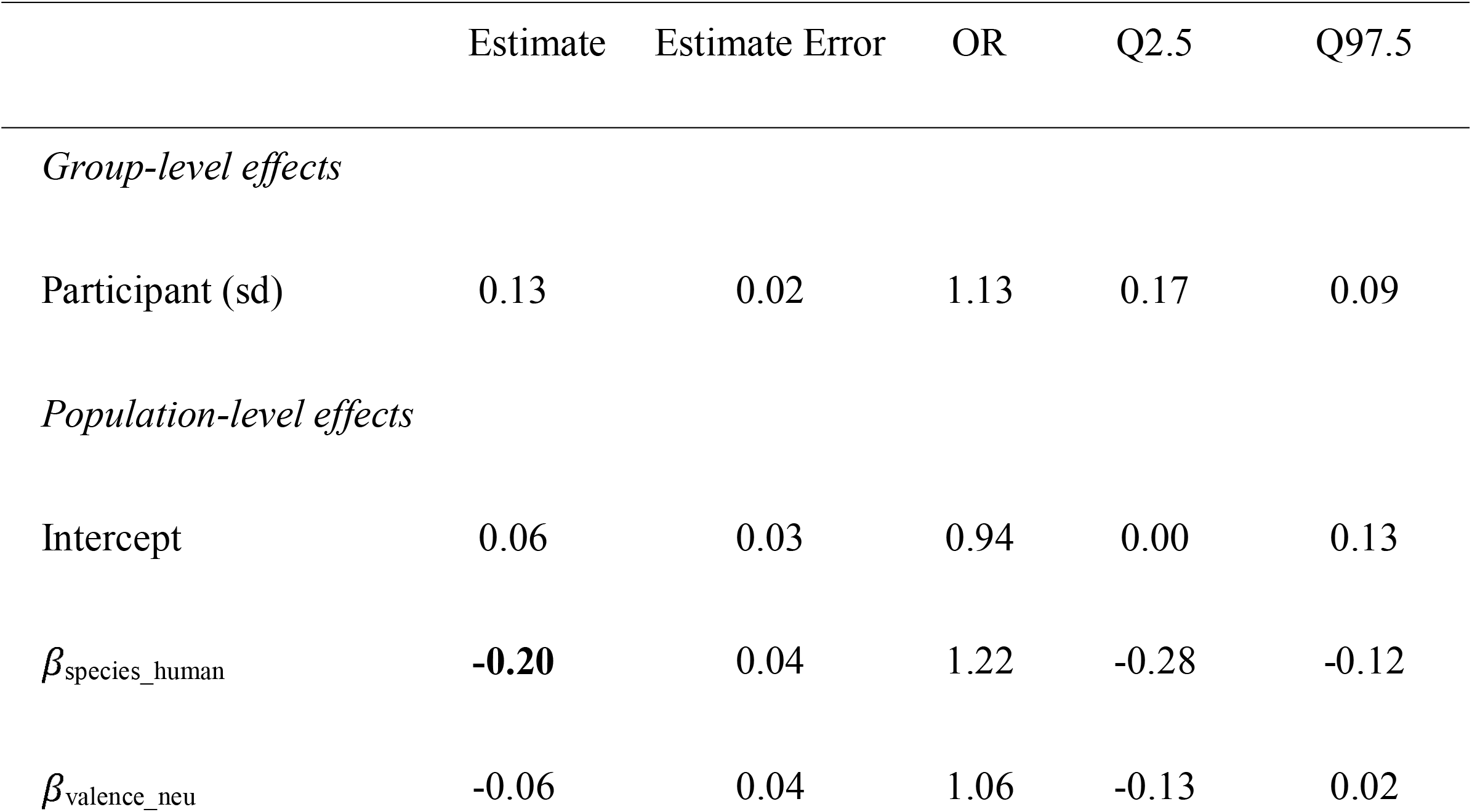
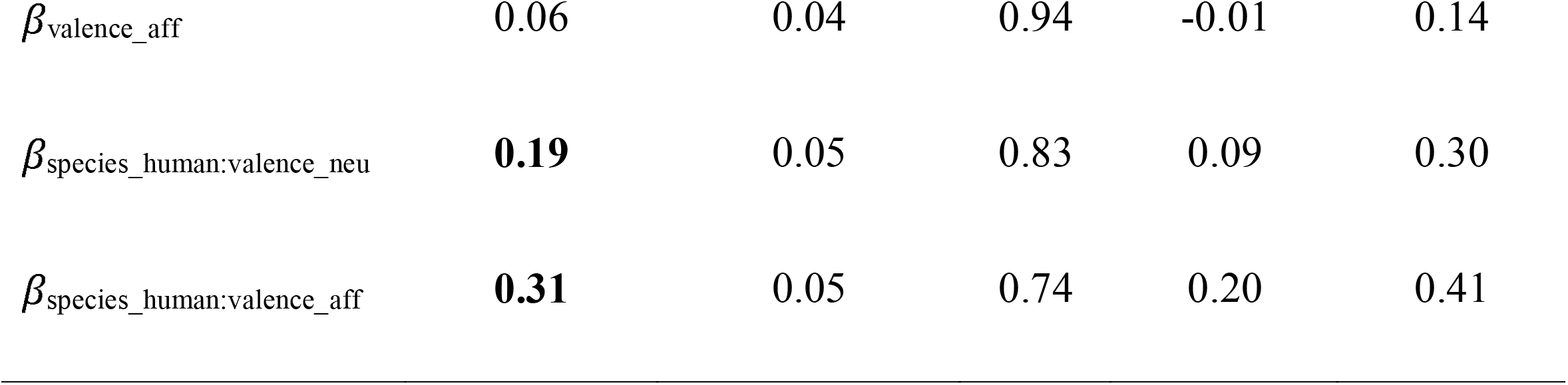
Logit transformed regression coefficients (posterior mean, standard error, 95% credible intervals and Bayes factor) and odds ratios (OR) of the beta distribution (positivity bias as a function of the two AOIs in the target screen). Reference category: Chimpanzee/agonistic.

|  | Estimate | Estimate Error | OR | Q2.5 | Q97.5 |
| --- | --- | --- | --- | --- | --- |
| *Group-level effects* |  |  |  |  |  |
| Participant (sd) | 0.13 | 0.02 | 1.13 | 0.17 | 0.09 |
| *Population-level effects* |  |  |  |  |  |
| Intercept | 0.06 | 0.03 | 0.94 | 0.00 | 0.13 |
| $\beta_{\text{species\_human}}$ | **-0.20** | 0.04 | 1.22 | -0.28 | -0.12 |
| $\beta_{\text{valence\_neu}}$ | -0.06 | 0.04 | 1.06 | -0.13 | 0.02 |
| $\beta_{\text{valence\_aff}}$ | 0.06 | 0.04 | 0.94 | -0.01 | 0.14 |
| $\beta_{\text{species\_human:valence\_neu}}$ | **0.19** | 0.05 | 0.83 | 0.09 | 0.30 |
| $\beta_{\text{species\_human:valence\_aff}}$ | **0.31** | 0.05 | 0.74 | 0.20 | 0.41 |

**Figure 2.**
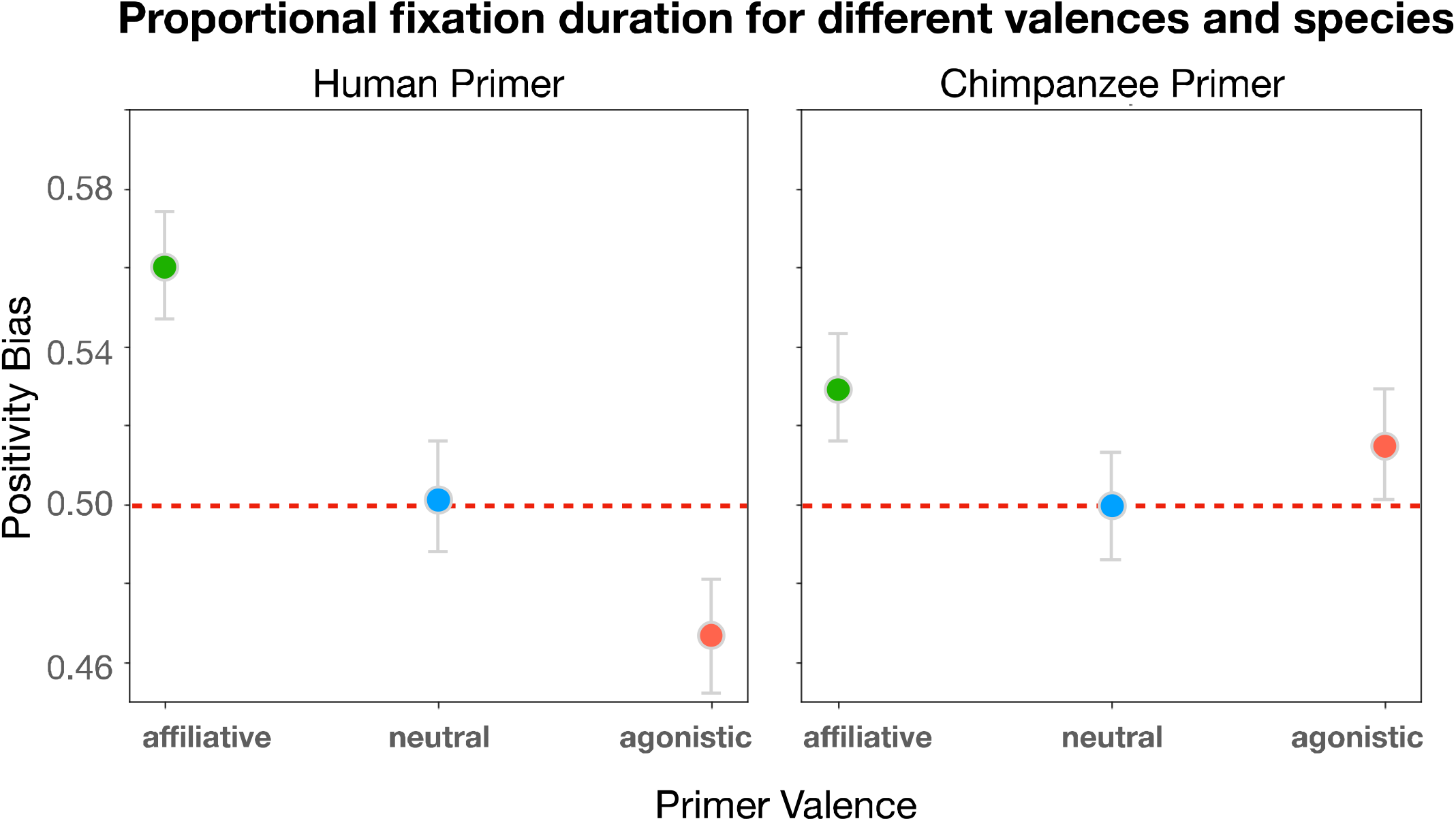
*Note*: Graphs displaying the proportional fixation duration (predicted model data) to emotional stimuli (positivity bias) of conspecifics and heterospecifics by human participants. Error bars reflect the 95% credible interval, dots represent the median. The measure of the positivity bias score higher than 0.5 indicates a longer fixation duration towards the positive emotional expression in the target screen. Hence, a positivity bias score lower than 0.5 indicates a longer fixation duration towards the negative emotional expression.

Investigating the interactions in the model (Table 2) reveals whether the strength of the positivity bias introduced by the valence effect differs for seeing primers of different species. For chimpanzee primers, no significant mean simple effect of valence was found. However, the tendency of positivity biases shifts shows that upon seeing affiliative stimuli, the bias increased (*β*_valence_aff_ = .06), meaning that positively valenced stimuli influence the participant to look more prevalently to the affiliative target picture. Contrary to our expectations, neutral chimpanzee primers compared to agonistic chimpanzee primers are followed by an decreased positivity biases towards the target pictures (*β*_valence_neu_= −.06). These simple effects are only reflecting tendencies and are not robust.

For the interactions comparing the simple effects of different valences between species, we found both effects to be significant. Comparing the neutrally valenced primers across species reveals that human neutral primers increased positivity bias more reliably than chimpanzee neutral primers (*β*_species_human:valence_neu_ = .19, *CI*[0.09 - 0.30], *OR* = 0.83). The main effect of neutrally human valenced primers is therefore *β*_H_valence_neu_= .13 (−.06 + .19 = .13). Similarly, comparing positively valenced primers across species yielded a significant interaction effect (*β*_species_human:valence_aff_ = .31, *CI*[0.20-0.41], *OR* = 0.74), showing that human affiliative primers reliably introduce positivity bias in the participants (compared to agonistic primers). The main effect on positivity bias upon seeing a positively valenced human primer is *β*_H_valence_aff_ = .37 (.06 + .31 = .37). An overview of the results is presented in Table 2 and Figure 2. The parameters of the beta distribution’s precision, the zero-one inflation and the conditional-one inflation are available in the Supplementary Material Table 1.

## Discussion

Understanding the emotions of others is a crucially valuable skill for social animals to successfully master group interactions and to navigate in their environment (de Waal, 2011). The ability to recognize and distinguish facial expressions that communicate affective states helps to predict the subsequent action of an encountering individual and can be highly impactful on an individual’s survival (Frijda, 2010). Since also the perception of other species’ emotional displays can influence the chances of survival (Jacobs & Vaske, 2019), we investigated to what extent emotion processing mechanisms in humans are transferable to accurately perceive and interpret the emotional expressions of our closest living relative, the chimpanzee. Introducing a prime-target paradigm, we compared how our participants’ attentional bias is influenced by emotional expressions of their own, as well as by the other species. As hypothesized, the attentional shifts occurred towards the targets that were congruent with the previously seen primer. Importantly, this effect was robust for humans viewing human emotional expressions, but partly contradictory and weaker for viewing chimpanzee emotional expressions. In our study, we confirmed that priming with emotional facial expressions introduces attentional biases toward emotional stimuli of conspecifics. Participants looked reliably longer at an emotional target stimulus that was congruent with the valence of the conspecific primer they saw before. Hence, a priorly presented affective stimulus depicting a human changed the amount of attention that participants allocated to afterward presented positive and negative visual information. Eventually, most likely due to their less pronounced relevance, neutral primers did not introduce this effect. The demonstrated priming effect elicited by positively and negatively connotated emotional displays confirms previous findings about the moderating role of valenced primers (Smith et al., 2006).

Contrary to our predictions, we did not find heterospecific (chimpanzee) primers to influence participants’ attention in a comparable way to conspecific (human) primers. This result is somewhat surprising, as previous research has demonstrated humans to be equally sensitive toward social cues from both species (Hattori, Kano, Tomonaga, 2010; Kret, Muramatsu & Matsuzawa, 2018). However, testing functional implications, in the present study we did not find the attentional bias of chimpanzees to have a robust priming effect. In regard to the negatively and neutrally valenced chimpanzee primers, the tendency of attentional bias shifts was somewhat comparable to the shifts upon viewing primers depicting humans. Positively valenced primers increased positivity bias, whereas neutral primers did barely change the fixation duration towards the two emotional targets. The negatively valenced primers did not have a robust impact on attention.

To verify that the participants’ correctly encoded chimpanzees’ facial expressions, we analyzed their valence and arousal ratings for the priming stimuli of both species. We found that negative emotional expressions were rated accordingly with more negative valence scores, as well as positive emotional expressions were rated accordingly with more positive valence scores. Ratings in valence mirrored the depicted emotional expressions’ categories, irrespective of the species. Similarly, we found emotional stimuli (positive and negative) to induce higher arousal in the participants, compared to neutral stimuli. These findings are in line with previous studies on the emotional perception of different species (Berlo, 2022; Kret & Matsuzawa, 2018) and confirm participants’ understanding and correct classification of chimpanzee emotional expressions. Furthermore, comparisons with ratings that were assigned to human primers show that the perceived valence and arousal of the emotional stimuli depicting different species are fairly similar. These data support a developed sensitivity for the perception and successful discrimination of emotional expressions in our close living relatives and possibly other related animals. This sensitivity might be contingent upon the extent of shared characteristics (Parr, Cohen & de Waal, 2005; Parr, Hopkins & de Waal, 1998). Arguably, close evolutionary relatedness is advantageous to demonstrate this skill, but not necessarily the only critical component. Humans being able to process and recognize e.g., dogs’ emotional states (Konok, Nagy & Miklósi, 2015) shows that the individual accumulated experience with another species may act as an additional covariate for understanding emotions in heterospecifics. We were able to validate the rating results of the participants with the rating results by experienced primate social cognition experts.

Given the participants’ validated understanding of chimpanzees’ emotional expressions, the induction of comparable effects on i.e., attention bias should conceivably be feasible. However, our results yielded a tendency of a positivity bias increase (negativity bias decrease) upon viewing a negatively valenced chimpanzee primer. Zooming in on potential explanations for this contradictory observation, the selected chimpanzee stimuli need to be closer examined. For the stimulus sub-set communicating negative affective states in chimpanzees, we chose one of the most frequently observed facial expressions across non-human primates: the bared-teeth display (Kim & Kret, 2022). Although in chimpanzees the expression predominantly occurs in agonistic interactions (Waller & Dunbar, 2005), it can signal different affective states in other species. Moreover, it is far from being one-dimensional, since morphological variants of the bared-teeth display have been found to signal different information and yield different social interaction outcomes (Clark et al., 2020, Martin-Malivel et al., 2007). Important in this context is that the activation of AUs that highly overlap with AUs forming a bared-teeth display, can resemble a smiling face in humans (Van Hooff, 1972; Parr & Waller, 2006). Hence, some participants of the present study might have misinterpreted the negative affective state in chimpanzees for a smile (perhaps due to canine visibility) and evaluated it as a positive expression, which might have averaged out the expected effect. Kret and van Berlo (2021) found an identical leakage effect from one emotional category to another in children who perceived pictures of distressed bonobos rather positively than negatively. The study argued that children, as opposed to adults, have not yet learned to take contextual information into account and incorporate this for their interpretation of an emotional expression. In the current eye-tracking study, this misclassification of the bared-teeth display occurred in the prime-target paradigm where the stimulus was presented on the screen for 2 seconds, but not in the post-hoc rating questionnaire where participants had no time restriction for indicating the perceived valence and arousal of the viewed primers. From this mismatch in the participant’s emotion classification abilities, we can conclude that the exposure time to an affective stimulus might play a consequential role. Supporting this relation, Lohse and Overgaard (2019) found perceptual awareness of faces to increase gradually with duration of the presented stimuli. Thus, future studies investigating humans’ understanding of other animals’ emotions should take the time exposure aspect into account and eventually increase the presentation duration of heterospecific primer pictures.

One caveat of the current study that should be explored further is the categorization of different emotional expressions. Establishing salient categories and classifying human expressions has been a great challenge, with disagreement in the field. Since expressions in non-human primates have been studied far less, the disagreement in their categorization is even more pronounced, introducing increased variability in experimental designs. Quantifying the neural, physiological, and phenomenological organization of emotions in humans has produced a categorical structure of various emotions across different sources for arousal and brain activity (Nummenmaa & Saarimäki, 2019). Similar neuroimaging experiments and multivariate pattern recognition analyses could be applied using chimpanzee emotional stimuli to incorporate the different domains of arousal for (proposed) distinct emotion categories. Comparing these activations for heterospecific and conspecific stimuli would potentially reveal homologies in emotion expression, perception, and processing in different species. In the present study, we chose two opposing valences to test the effects of priming on attention, however, even with ecologically validated stimuli, we presumably encountered partial leaking from one category to another. Studying a larger variety of emotional expressions in non-human primates might arguably first increase such overlaps of categories but in the long-term help to distinguish salient cues and benefit the more precise identification of facial expressions. Possibly, instead of the distinct categorization into exclusive categories, future research could follow the approach of assigning animal emotions on a continuum, exploring dynamic facial movements and their communicative signals (Jack, Garrod & Schyns, 2014). Further regarding stimuli selection, follow-up studies could investigate affective state induction using dynamic stimuli. In the current study, we used static pictures of facial expressions in humans and chimpanzees which might have limited the participants’ ability to recognize an affective state correctly. Previous investigations have shown that moving faces are generally better recognized compared to still faces (Knight & Johnson, 1997) since a dynamic scene provides more communicative signals and includes more contextual cues. Overall, an increased amount of information facilitates the perception of an emotional expression (Parr, 2003; Seyfarth, Cheney & Marler, 1980) and hence might induce a stronger priming effect (Garrido-Vásquez et al. (2018).

In addition to the methodological concern about selecting stimuli depicting equivalent facial expressions in different species, our study faces another constraint regarding the match between chimpanzee and human primers. While all chimpanzee photographs were taken opportunistically in a naturalistic setting, the human photographs as a subset of the Chicago Face Database were showing acted emotions in a highly standardized environment. This introduces an additional disbalance to the comparison of the presented stimuli and limits the experimental control, which might have influenced the perception of emotions, as previous studies have demonstrated (Hideg & van Kleef, 2017). While it would have been feasible to incorporate human stimuli from naturalistic contexts, we deliberately abstained from doing so due to its limited exploration within the current body of literature. Our intention was to extend upon prior research efforts. Tackling the standardization problem from another side, recent studies have started creating morphologically altered, hyper-realistic facial expressions in non-human primates for studies on visual processing and social cognition (Murphy & Leopold, 2019; Wilson et al., 2019). Using advanced high-tech software programs, some teams have been able to produce highly realistic scenes depicting various facial and body movements in monkeys and great apes, overcoming the uncanny valley (Siebert et al., 2020; Kawaguchi, 2021).

Based on the parallels between humans and non-human primates that since Darwin’s pioneering work have only unfolded further, it can be assumed that there is an evolutionary continuity in the emotional behaviors and their processing in humans and non-human primates (Kim & Kret, 2019; Kret et al., 2020). Aiming to find homologies in human processing of facial expressions in their own species, and in an evolutionary closely related species, we tested the influences on attention introduced by priming with differently valenced emotional stimuli depicting humans and chimpanzees. Attention was shifted toward the emotional target picture that was congruent with the valence of the conspecific emotional expression depicted in the primer picture. Contrary to our expectations, we did not find this effect to occur with chimpanzee primers. Additional cross-species systematic investigations with slight adjustments are needed to fully address the gap of a shared evolutionary ancestry, and ultimately rule out the idea of emotions being unique to humans.

## Acknowledgments

We would like to thank Roy de Kleijn, Tom Wilderjans, Elio Sjak-Shie, Lukas Kunz, Alessandro Palumbo, Juan Olvido Perea-García, Evert Dekker, Yena Kim, and Tristan Matsulevits for their valuable feedback and discussions on the methods used in this work. This research is supported by the European Research Council (ERC) (Starting Grant #804582) awarded to M.E.K.

