## Supplementary Material for "Priming using Human and Chimpanzee Expressions of Emotion Biases Attention toward Positive Emotions"

I. **Validation of the Stimulus Material**

After the eye-tracking experiment, participants rated each primer (108 pictures) on two separate sliders. The dependent variables *emotional valence* and *arousal* (answers coded on sliders scaling from 0-100) were analyzed for the expert and the non-expert group using four analyses of variances (ANOVAs) in total. We investigated the differences in valence and arousal levels between primer species and emotional categories:1) valence ratings for different emotional categories and primer species, 2) arousal ratings for different different emotional categories and primer species, 3) valence rating comparison between experts and non-experts, 4) arousal valence rating comparison between experts and non-experts. A general overview of the rating distributions for the two dependent variables can be found in Figure 1A and 1B.

First, we found the valence ratings to significantly differ between the emotional categories of the primer pictures (*F*(2, 98) = 431.54, *p* <.05), meaning that the participants robustly indicated different valences to primers of different emotional categories. From the outcome scores, we can extract that negative valenced primers were assigned the lowest values, neutral primers scored in the middle, and positive primers were assigned the highest values on a slider ranging from the negative to the positive category. Which species was depicted on the primer picture did not influence these valence ratings significantly (*F*(1, 49) = 3.21, *p* =.01), however, the interaction effect between *species* and *emotional category* was significant (*F*(2, 98) = 56.45, *p* <.05). Similarly, the same pattern applies when comparing the ratings of the negative with the positive category for both species. For the negative primer category compared with the neutral category, the difference in ratings was the same for chimpanzee and human primers.

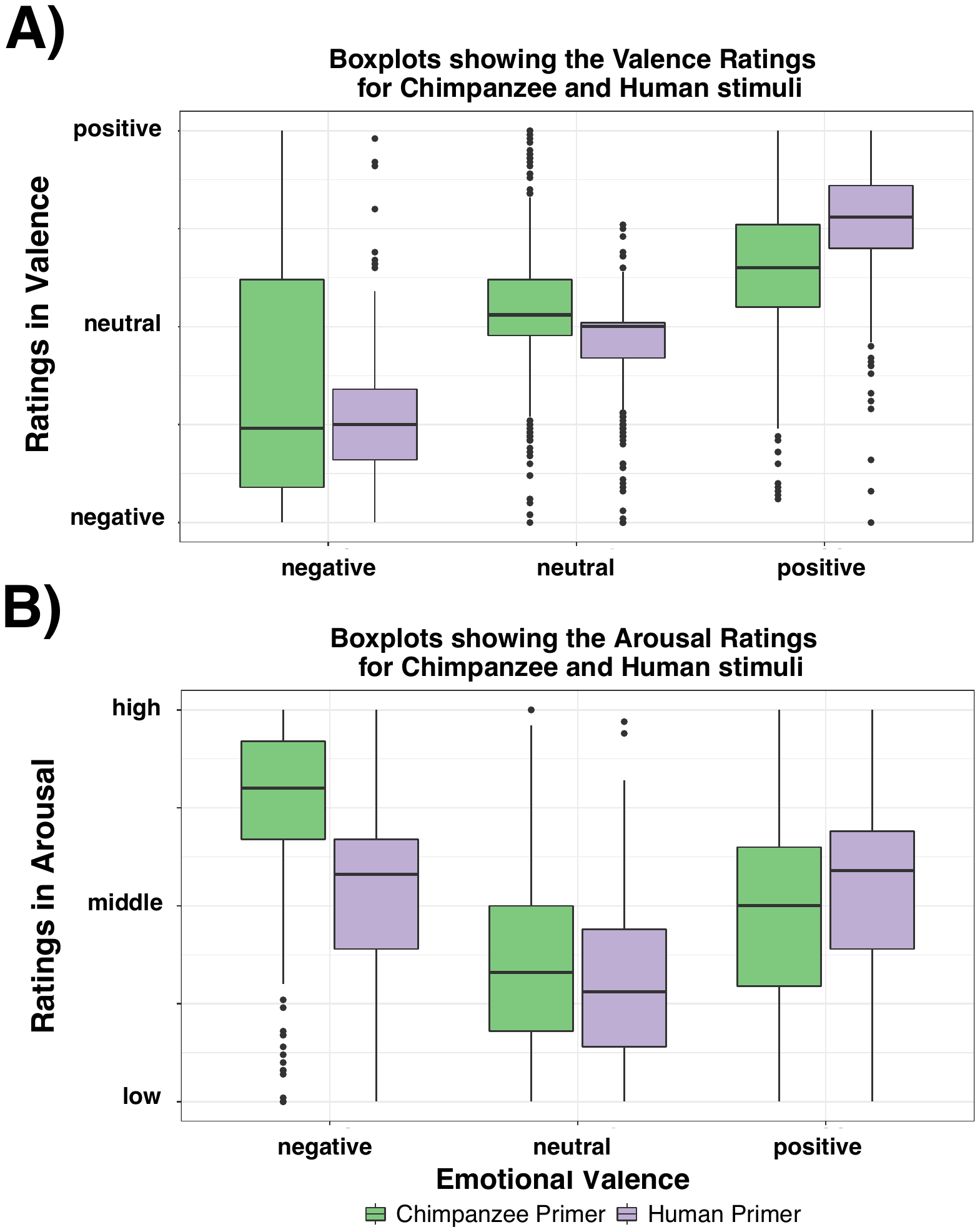
**Figure 1**

*Note*. Boxplots visualizing the distribution of the valence and arousal ratings for each emotional category, separated by species. Error bars reflect the interquartile range (Q3-Q1), dots represent the outliers.

We then analyzed the arousal levels elicited by the primer pictures of different emotional categories depicted in different species. We did find significant differences in the arousal ratings for the three *emotion categories* positive, negative and neutral (*F*(2, 96) = 180.11, *p* <.05). From the outcome scores we can extract that in general, negatively and positively valenced primers elicited higher arousal in the participants compared to neutral primers (Figure 1B). Furthermore, the main effect of *species* was also significant with *F*(1, 48) = 40.06, *p* <.05). This means, that the arousal level ratings differed between seeing a chimpanzee primer and seeing a human primer. Overall, arousal levels for chimpanzee primers were indicated to be higher, except in the *emotion category* “positive”. We also found an interaction effect between *emotion category* and *species* to be significant (*F*(2, 96) = 126.60, *p* <.05). As previously observed in the valence ratings, the differences in the arousal ratings for the emotion categories “neutral” and “positive” differ between the depicted primer species. Arousal looking at positively valenced pictures was indicated to be higher for human facial expressions compared to chimpanzee facial expressions. For negatively valenced primer pictures, an opposite pattern appeared: ratings indicated higher arousal for negative chimpanzee emotional expressions compared to negative human emotional expressions.

Lastly, in order to validate whether the chimpanzee stimuli were correctly classified by our participants, we compared their valence and arousal ratings to seven experts’ ratings of the same stimulus set. We did not find a significant difference for valence ratings between experts and non-experts (*F*(1, 49) = 0.96, *p* = 0.33), meaning that the participants of the study, who were not experienced with non-human primates’ emotions did not classify chimpanzee emotional expressions differently from non-human primate experts (Supp. Figure 2 A). This confirms the participants’ understanding of the emotional stimuli in the other species and validates our stimulus selection. Furthermore, we did not find differences in the indicated arousal levels between experts and non-experts (*F*(1, 48) = 0.44, *p* = 0.51) (Supp. Figure 2 B). From these results we can conclude, that the selected primers were perceived similarly irrespective of the expertise and experience with emotional expressions in chimpanzees.

**Figure 2**
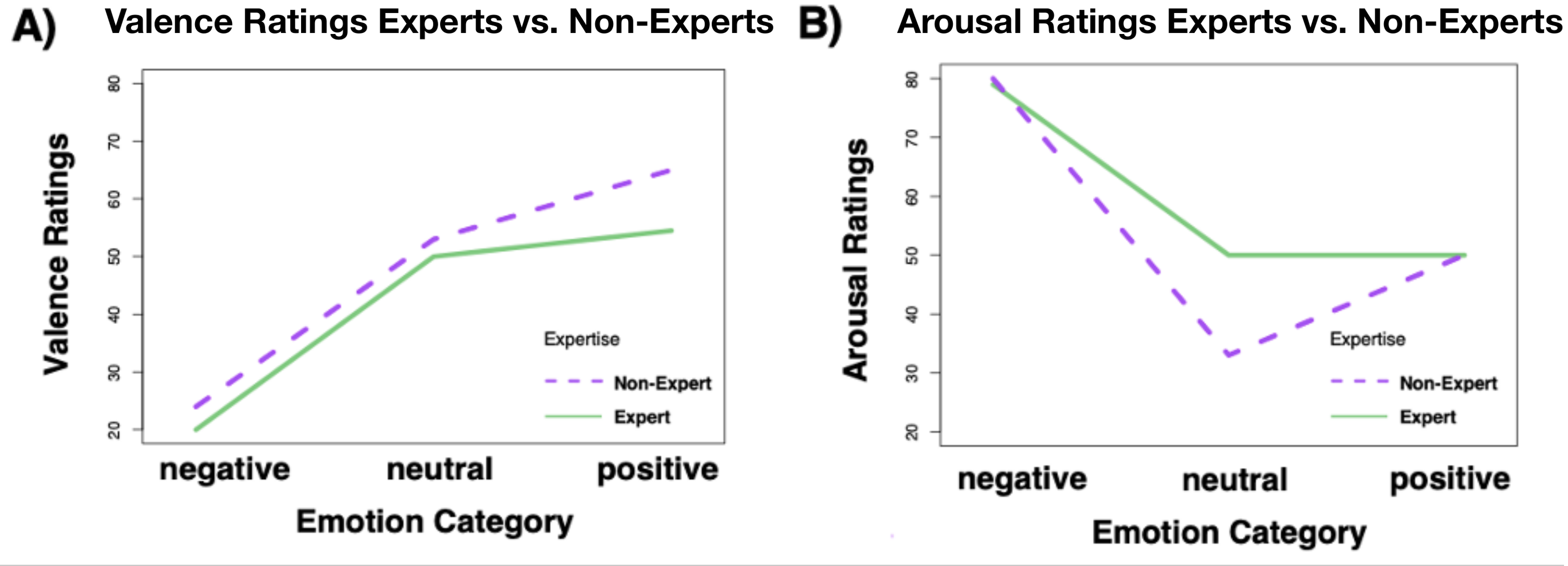

*Note*. Graphs visualizing the differences between experts’ and non-experts’ ratings of chimpanzee emotional expressions in A) valence for the different emotion categories, and B) arousal for the different emotion categories.

II. **Information on the model notation**

Model notation for the brms model, implementing the zero–one-inflated beta model and the prior distribution via R.

form <- bf(bias_score ~ P_SPECIES * P_VALENCE + (1|Participant),

phi ~ P_SPECIES * P_VALENCE + (1|Participant),

zoi ~ P_SPECIES * P_VALENCE + (1|Participant),

coi ~ P_SPECIES * P_VALENCE + (1|Participant))

prior_list = c(prior(normal(0, 0.25), class="Intercept"),

prior(normal(0, 0.5), class="b"),

prior(normal(0, 0.5), class="b", dpar="coi"),

prior(normal(0, 0.5), class="b", dpar="zoi"),

prior(normal(0, 0.5), class="b", dpar="phi"))

model <-brm(form,

data = targetscreen_data_cleaned, iter = 6000, warmup = 1000,

cores = 2, control=list(adapt_delta=0.99),

prior = prior_list,

family=zero_one_inflated_beta())

To compare the fixation duration of the AOIs in the target screen, we estimated the fixation proportion (positivity bias) using a Bayesian zero–one-inflated beta multilevel model. The model considers a beta distribution for the continuous proportion outcome in the closed (0,1) interval. Model coefficients of the beta distribution’s precision (phi) on the scale of the log-link function, the zero–one inflation (zoi) on the logit scale and the conditional one inflation (coi) on the logit scale for the calculated negativity bias (proportional fixation duration at the negative AOI).

**Table 1**

|  | | Estimate | Estimate Error | Q2.5 | Q97.5 |
| --- | --- | --- | --- | --- | --- |
|  | Phi_Intercept | 1.60 | 0.08 | 1.44 | 1.77 |
|  | Zoi_Intercept | -3.47 | 0.27 | -4.01 | -2.94 |
|  | Coi_Intercept | -0.11 | 0.30 | -0.72 | 0.48 |
|  | Phi_SPECIES_H | 0.08 | 0.06 | -0.04 | 0.20 |
|  | Phi_VALENCE_NEU | 0.05 | 0.06 | -0.07 | 0.18 |
|  | Phi_VALENCE_POS | 0.01 | 0.06 | -0.12 | 0.13 |
|  | Phi_SPECIES_H*VALENCE_NEU | -0.22 | 0.09 | -0.39 | -0.05 |
|  | Phi_SPECIES_H*VALENCE_POS | -0.15 | 0.09 | -0.32 | 0.02 |
|  | Zoi_SPECIES_H | 0.26 | 0.18 | -0.09 | 0.62 |
|  | Zoi_VALENCE_NEU | 0.35 | 0.19 | -0.03 | 0.74 |
|  | Zoi_VALENCE_POS | 0.15 | 0.19 | -0.22 | 0.52 |
|  | Zoi_SPECIES_H*VALENCE_NEU | -0.75 | 0.25 | -1.24 | -0.26 |
|  | Zoi_SPECIES_H*VALENCE_POS | -0.12 | 0.26 | -0.62 | 0.38 |
|  | Coi_SPECIES_H | -0.07 | 0.26 | -0.59 | 0.44 |
|  | Coi_VALENCE_NEU | -0.23 | 0.29 | -0.80 | 0.33 |
|  | Coi_VALENCE_POS | -0.06 | 0.28 | -0.62 | 0.50 |
|  | Coi_SPECIES_H*VALENCE_NEU | 0.43 | 0.34 | -0.25 | 1.10 |
|  | Coi_SPECIES_H*VALENCE_POS | 0.57 | 0.35 | -0.12 | 1.25 |
